# Re-inventing sex: A W chromosome and a path out of parthenogenesis for Bunonematidae sp. nematodes

**DOI:** 10.64898/2026.07.28.741199

**Authors:** Janna L. Fierst, Lara Arbex Herden, Christina A. Burns, Victoria K. Eggers, Pushpalata Kayastha, Michelle A. McCauley, Jasbelle Sosa, Karolina Willicott

**Affiliations:** Biomolecular Sciences Institute, Florida International University; Department of Biological Sciences, Florida International University, 11200 SW 8th Street, Miami, 33199, Florida, USA

**Keywords:** evolution, genetics, sex chromosomes, nematodes, contingency

## Abstract

Separate sexes are a conserved character across the tree of life but the molecular and genetic underpinnings of sex determination are remarkably diverse. Here we describe the evolution of a novel sex chromosome system in the nematode Bunonematidae sp. JU3390. Combining deep PacBio long read sequencing, HiC chromatin capture and Illumina sequencing of individual male and female Bunonematidae sp. JU3390 we identified an autosome-derived genomic segment currently functioning as a female-limited W chromosome. Phylogenetic and orthologous gene analyses suggest this species passed through a period of parthenogenesis before transitioning back to male-female outcrossing. All other described nematodes have male heterogametic (XX/X0 or XX/XY) chromosome systems, suggesting that in re-evolving sex Bunonematidae sp. JU3390 also evolved novel female heterogamety. We identified large structural deletion mutations segregating in the population including a 988kb portion of Scaffold 5 that, if homozygous, would result in lethality. We propose this neo sex chromosome system evolved to escape lethal structural mutations accumulated during a period of parthenogenesis, demonstrating the role of evolutionary contingency in this rapid re-invention of sex.

## INTRODUCTION

Across the tree of life there are two dominant sex chromosome systems^1,2^. In XY systems females are homogametic carrying two X chromosomes while males are heterogametic, carrying one X and one Y. In ZW systems males are the homogametic sex carrying two Z chromosomes while females are ZW heterogametes. Although there is broad organismal diversity in sex chromosomes these systems have, to date, been phylogenetically limited^3^. For example, mammals and insects have XY systems while birds, snakes and butterflies have ZW systems^1^. Sex chromosomes define development but their evolutionary dynamics remain a paradox; they are deeply conserved in some clades yet remarkably labile in others^4,5^. Chromosome systems impact multiple dimensions of organismal evolution including dosage compensation, gene regulation, development and overall patterns of mutation including dominance dynamics, Faster X evolution^6^ and Haldane’s rule^7^, that the heterogametic sex is vulnerable in hybridization and speciation. Sex chromosome transitions have been reported in cichlid fish^8^, frogs^9^ and willows^10^ but the evolutionary conditions that select for XY or ZW systems, transitions or stability are not understood.

In nematodes females carry two X chromosomes with males carrying either a single X (X0)^11^ or in rare cases a male-specific Y^12–14^. This system is deeply conserved with nematodes in the Rhabditina group (Figure 1A) showing conservation of ancestral chromosomal identity across both autosomes and sex chromosomes^15^. Analysis of orthologous genes has identified conserved ’Nigon elements’ capable of tracking chromosomal identity across this group, representing over 350 million years of evolution^16^. Exceptions to Nigon conservation and chromosomal identity have been reported in nematodes with non-Mendelian segregation^17^ and parthenogenetic species where it’s thought that asexual reproduction provides a release from the strict genetic constraints of homologous crossover and recombination^18^. For example, the plant parasitic nematode *Meloidogyne-hapla* shows low Nigon conservation and reproduces through facultative meiotic parthenogenesis, regularly producing selfed offspring and infrequently outcrossing with migratory males^19^.

**Figure 1:**
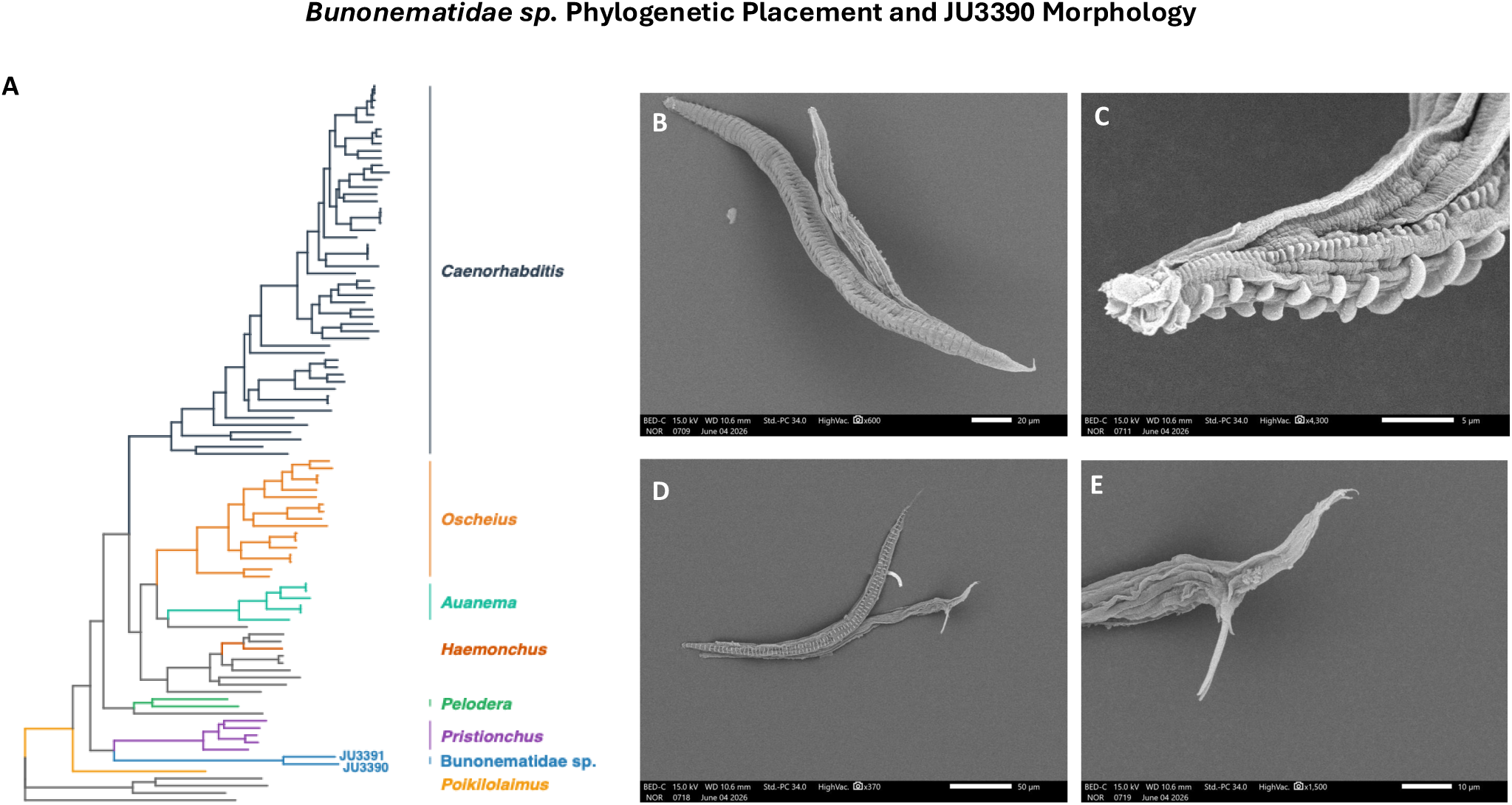
Bunonematidae sp. JU3390 and JU3391 are at the base of the Rhabditina group with males readily identified in Bunonematidae sp. JU3390. A) Phylogenetic reconstruction placed Bunonematidae sp. JU3390 and JU3391 at the base of the Rhabditina group. B) Electron microscopy of Bunonematidae sp. JU3390 demonstrated the characteristic morphology, unusual among Rhabditina with lateral lines and tubercules, noticeable in C) close-up imaging. D-E) Male spicules were readily identified in desiccated samples.

Reproductive mode diversity is common in nematodes. The *Meloidogyne* group contains a spectrum of reproductive strategies ranging from obligate outcrossing and meiosis to strict mitotic parthenogenesis^20^. Many species including the model self-fertile nematode *Caenorhabditis elegans* maintain the ability to outcross with rare males, theoretically benefiting from both the stability and offspring production of self-fertility and the evolutionary ability to purge mutation and create new genetic combinations. A long-standing question for evolutionary biology is, however, the persistence of multiple parthenogenetic species^21^. According to theory these species represent evolutionary dead ends^22^. Losing the ability to purge deleterious mutations creates a continually ticking ‘ratchet’ where build-up of deleterious variation eventually leads to species extinction^23–25^. How species may escape this process is unclear as few natural examples have been identified.

Here we describe the evolution of a novel female heterogametic ZW sex chromosome system in Bunonematidae sp. We used a combination of deep Hifi sequencing and HiC chromatin capture to reconstruct chromosomes for two bunonematid worms, JU3390 and JU3391. Using phylogenetic reconstruction and orthology analyses we found these Bunonematidae sp. have poor Nigon conservation, suggesting an evolutionary history of parthenogenesis (Supplementary Figure 1). One species, Bunonematidae sp. JU3391, has only females and minimal rates of molecular variation consistent with automictic parthenogenesis. A second species, Bunonematidae sp. JU3390, outcrosses regularly with populations split between males and females^26^ (Figure 1B). Using single worm DNA sequencing we identified a sex-associated chromosomal fragment in JU3390, female-limited and functioning as a ‘W’ chromosome. We also identified multiple large segregating structural mutations in Bunonematidae sp. JU3390 including a 988kb region on Scaffold 5 that, if homozygous, would lack essential genes. Together, these results suggest Bunonematidae sp. JU3390 passed through a period of partial or facultative parthenogenesis, accumulating deleterious variation through a structural ‘Muller’s ratchet’^21,23^. Population persistence depended on the maintenance of heterozygous variation, readily achieved through outcrossing, sex-capture of an autosome and a re-invention of sexual reproduction.

## RESULTS AND DISCUSSION

### A novel chromosome complex in Bunonematidae sp. JU3390

Bunonematidae is a Family of small, free-living worms at the base of the Rhabditina group (Figure 1A). The group has not been well-studied and is characterized largely by morphological taxonomic descriptions^27^. As we can not differentiate between genera within the group we refer to these nematodes as Bunonematidae sp. We used Pacbio Hifi to sequence pooled populations of Bunonematidae sp. JU3390 at high depth, approximately 341x coverage of the estimated 60Mb genome (Supplementary Table 1). Similarly, we sequenced pooled populations of Bunonematidae sp. JU3391 at approximately 339x coverage of the estimated 57Mb genome. For each species Hifiasm assembly with HiC libraries resulted in six contiguous assembled chromosomes (Figure 2A), echoing the conserved six chromosome karyotype shared across Rhabditina. The consensus diploid and individual haploid Bunonematidae sp. JU3391 assembled sequences were more than 99% identical with remarkably low molecular diversity of just 0.003% (1,738 single nucleotide variants; Supplementary Figure 1). However, we identified substantial polymorphism in the Bunonematidae sp. JU3390 individual assembled sequences.

**Figure 2:**
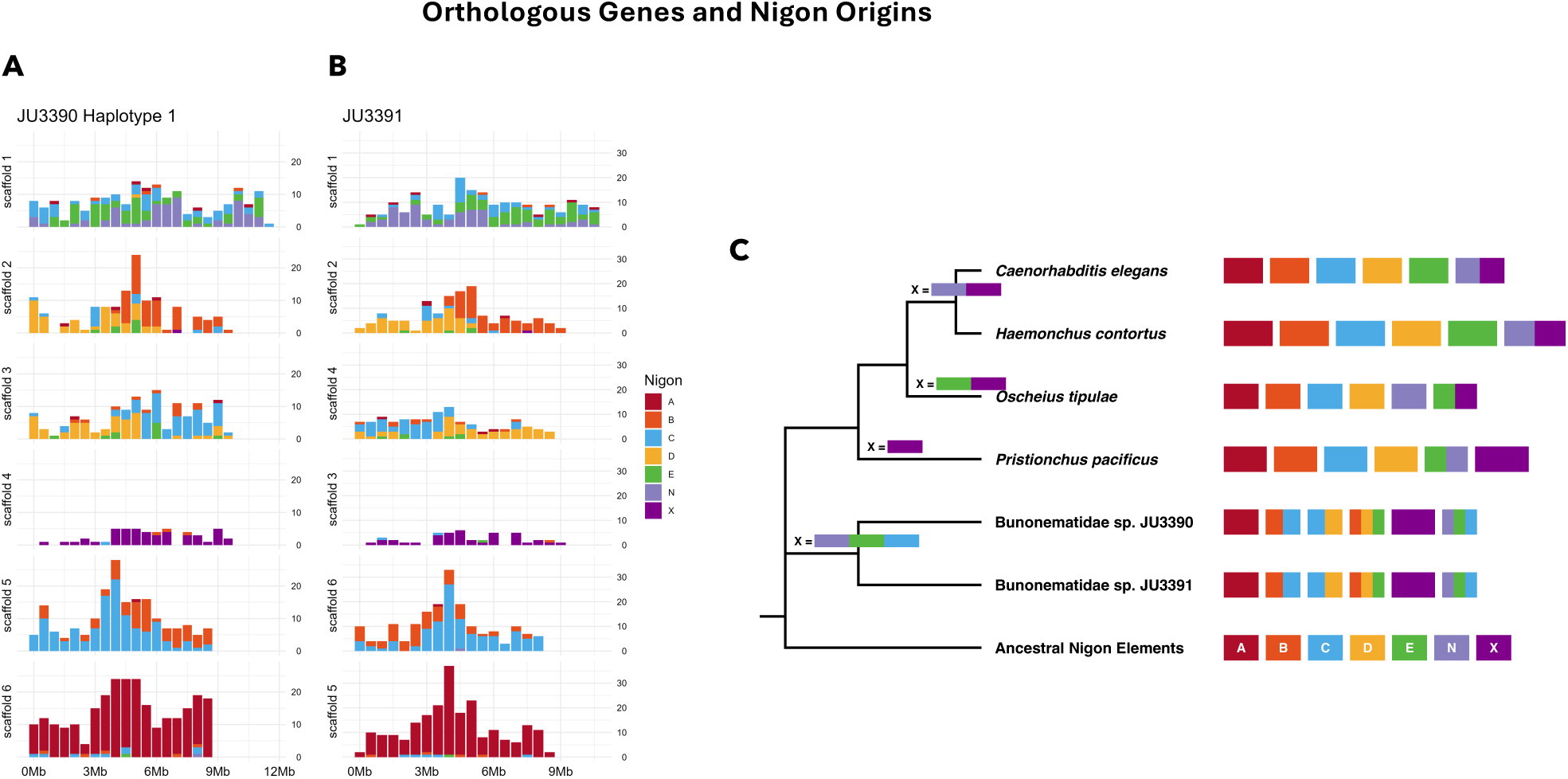
Assembled genome sequences for JU3390 produced 3 Scaffold 1 variants with high homology. A) The HiC contact map for Haplotype 1 showed 6 chromosomes. In B) Haplotype 2 and C) Haplotype 3 the Scaffold 1 sequence was absent and two smaller variants (*X_A_* and *X_B_*) were present. The HiC contacts between these fragments suggested D) these variants were fused (*X_neo_*) in a head-to-tail orientation in haplotype 2 and tail-to-tail in haplotype 3. E) Alignment of these Scaffold 1 / X variants demonstrated that *X_A_* and *X_B_* were fragments of the larger Scaffold 1. *X_A_* had shared colinearity with Scaffold 1 and roughly 4Mb of *X_B_* while *X_B_* had a 3.6Mb section that was inverted relative to Scaffold 1. Both SNVs and SVs were accumulated within the boundaries of this inverted region on Scaffold 1 and *X_B_*. In contrast very few SNVs or SVs were identified on *X_A_*.

In Bunonematidae sp. JU3390 the largest 11Mb Scaffold 1 in Haplotype 1 (Figure 2A) was not identified in Haplotypes 2 and 3. Instead, we identified extreme structural polymorphisms related to the Scaffold 1 backbone^28^. We found 8.9Mb *X_A_* and 7.8Mb *X_B_* fragments (Figure 2B-C) assembled in head-to-tail orientation in Haplotype 2 and tail-to-tail orientation in Haplotype 3 (Figure 2D). HiC contact maps indicated that *X_A_* and *X_B_* were physically fused into one *X_neo_* chromosome^29,30^. Alignment of Scaffold 1, *X_A_* and *X_B_* identified the two *X* variants as fragments of the larger ancestral Scaffold 1 backbone (Figure 2E). Analysis of segregating polymorphisms identified substantial numbers of single nucleotide variants (SNVs) and structural variants (SVs) accumulated at the alignment boundaries between Scaffold 1 and a highly contiguous but inverted portion of *X_B_.* The *X_A_* fragment showed high homology and 99.45% nucleotide identity with both Scaffold 1 and *X_B_*, suggesting these fragments are actively recombining. Ancestral origins and ultra high homology between these segments made it difficult to diagnose the karyotype from sequencing data, stressing the critical value of our high-resolution, high-depth HiC approach (Supplementary Table 2).

### Nigon element degradation and a history of parthenogenesis

The ancestral Rhabditina karyotype encompassed seven chromosomes, six autosomes and an ancestral X^15,16^. In current Rhabditina species the X chromosome is readily identified through orthologous genes, sex-specific segregation and size, with X typically larger than autosomes. In both Bunonematidae sp. JU3390 and JU3391 orthologous genes that originated from the ancestral Nigon X were located on Scaffold 4 (Figure 3A-B). This chromosome did not show sex-specific patterns and was smaller than other chromosomes, suggesting its role as a sex chromosome was retired in the past and its function is now autosomal. Scaffolds 4 and 6 contained genes originating largely from Nigon X and Nigon A respectively. The remaining scaffolds mapped to multiple Nigon origins, suggesting a complex history of chromosomal fusions and fissions. Scaffold 1, *X_A_* and *X_B_* orthologous genes mapped to Nigon C, E and N origins (Figure 3A; Supplementary Figure 2A-C). In comparison, Rhabditina worms show largely intact Nigon conservation (Figure 3C) and Nigon X inheritance. Across the group Nigon X orthologous genes are found in the present-day X chromosome in *Pristionchus pacificus* ^13^, fused with Nigon E in *Oscheius tipulae* ^16^ and fused with Nigon N at the base of the clade containing *C. elegans* ^17,31^. In each of these species Nigon X orthologous genes are found in the present-day sex chromosome, segregating as either XX/XO^11^ or in rare cases XX/XY^13,14^.

**Figure 3:**
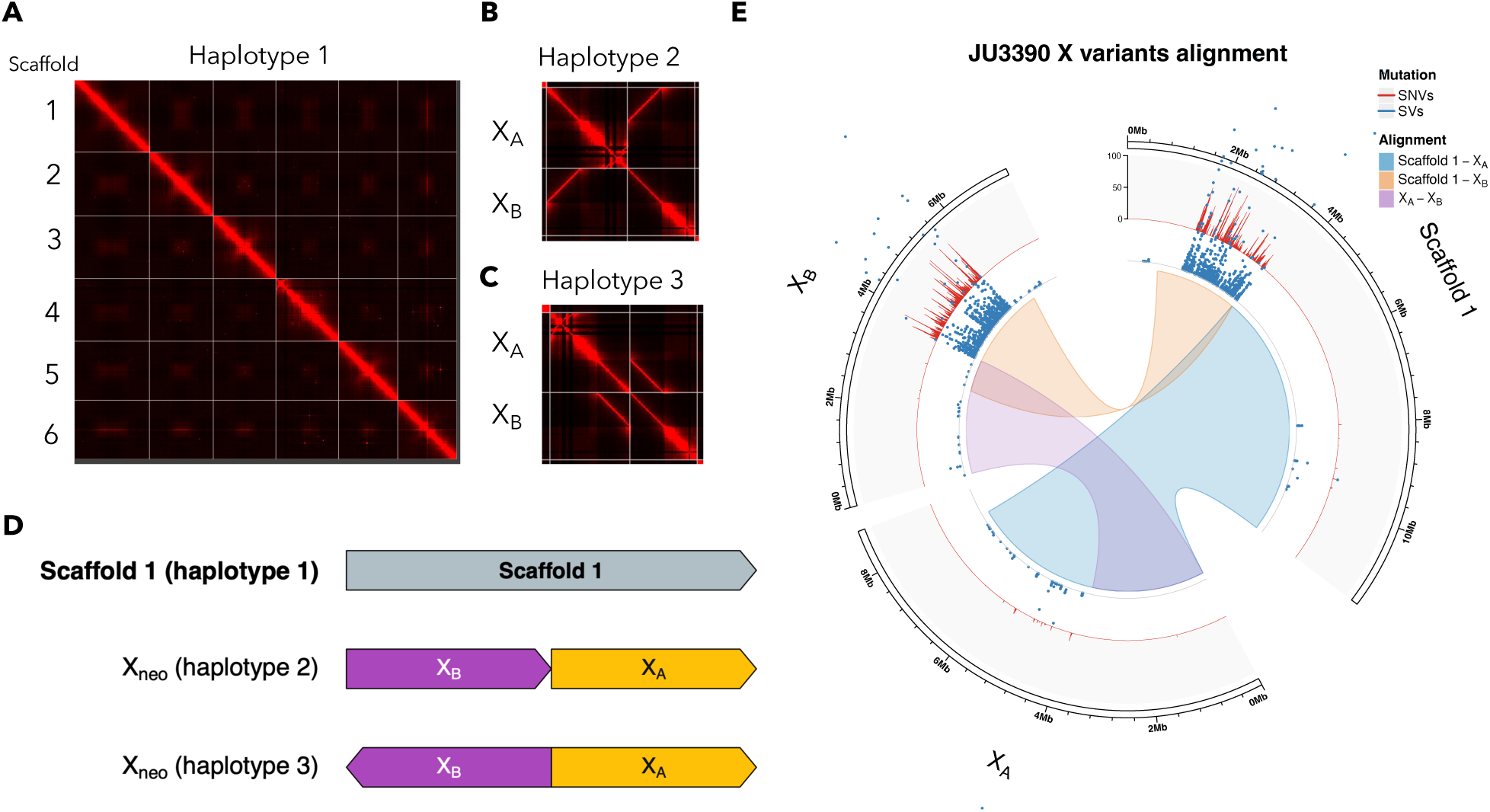
Bunonematidae sp. JU3390 and JU3391 are unique in both the mixture of Nigon elements found in their chromosomes and the Nigon C, E and N composition of their current sex chromosome. A) Identification of single copy orthologous genes showed that Bunonematidae sp. JU3390 and B) JU3391 share similar patterns of Nigon element origins across assembled scaffolds. Scaffold 1 and the X variants *X_A_* and *X_B_* contain genes from multiple Nigon elements with the majority originating in C, E and N. Scaffolds 1, 2, 3 and 5 contain genes from multiple Nigon elements indicating a history of chromosomal fusions and fissions. C) Rhabditine worms had 7 ancestral chromosomes and other species retained both Nigon identity in current karyotypes and Nigon X contribution to the modern X chromosome. Chromosome size is plotted on a *log*_10_ scale for visualization.

### Single worm sequencing identified a female-limited chromosome

Individual multiple displacement amplification (MDA) sequencing of 11 females and 12 males identified 4 distinct X variant karyotypes. A concordance score, combining breakpoint analysis with segment sequencing coverage, identified the presence of the 11Mb Scaffold 1 backbone in all sequenced individuals (Figure 4A). The *X_A_* segment was identified in all individuals with the exception of one male. High Scaffold 1 sequencing coverage suggested this individual carried two Scaffold 1 sequences. Two males had extremely high *X_A_* sequencing coverage, suggesting multiple segments in their karyotype (Figure 4B). The *X_B_* segment was identified in 9 of 11 females and absent in 9 of 12 males. Two males carried *X_B_* segments although high mapping variance and poor alignment statistics indicated these are deeply mutated and potentially rearranged. Two females carried no *X_B_* sequences and one male carried Scaffold 1, *X_A_* and *X_B_* segments. Bunonematidae sp. JU3390 are extremely small and difficult to sex without destroying the worm and we marked these discordant samples ‘Putative Females’ and ‘Putative Males’. Currently, we cannot definitively distinguish between misidentification or incomplete penetrance of chromosomal sex determination as environmental^32^, thermosensitive^13^ and stochastic sex determination^33^ have been reported in related worms. Collectively the competent female karyotype in Bunonematidae sp. JU3390 was Scaffold 1 / *X_neo_* while males carried multiple karyotypes, each lacking a contiguous, competent *X_B_* segment (Figure 4B; Supplementary Figure 3). These noisy segregation and structural mutation dynamics suggest this sex chromosome system is evolutionarily young and potentially in current transition^34^.

**Figure 4:**
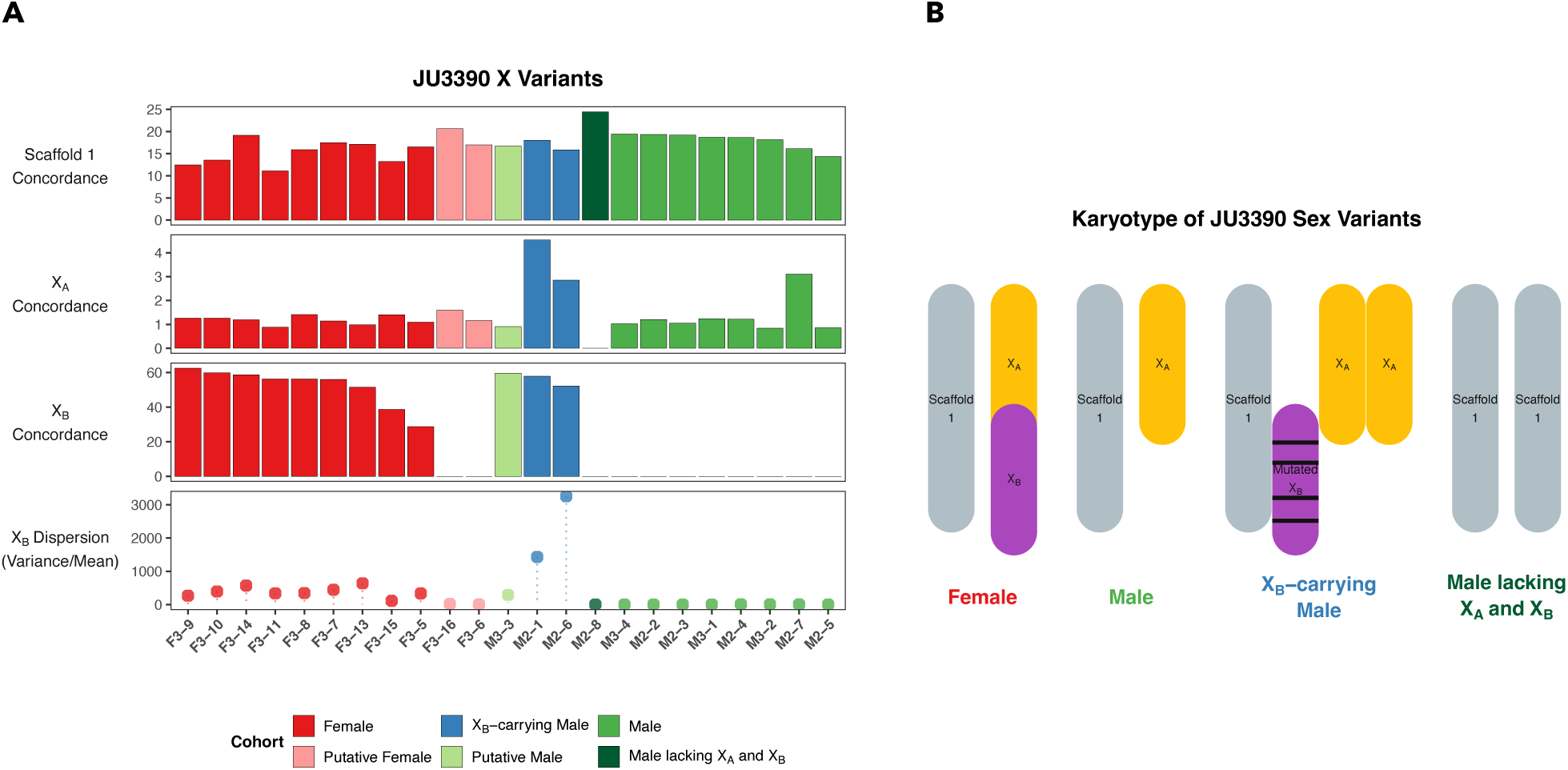
Single worm sequencing identified three karyotypes found in male Bunonematidae sp. JU3390 and one karyotype found in females. A) Alignment of Illumina MDA DNA sequence libraries combined with breakpoint analysis showed that all individuals contained Scaffold 1 with one male having high sequence read coverage, suggesting the presence of two Scaffold 1 fragments. *X_A_* was identified in 22 of 23 individuals and high sequence read coverage suggested multiple *X_A_* segments in two males. *X_B_* was identified in 9 of 11 females, suggesting either mis-sexing in the laboratory (putative females) or a dynamic relationship between X variants and phenotypic sex. *X_B_* was identified in 3 males although 2 of these individuals showed extremely high disperson and mapping variance suggesting a highly mutated, rearranged fragment. Nine males lacked the *X_B_* segment. B) The sequence read coverage and breakpoint analysis data suggested four potential karyotypes with females Scaffold 1 / *X_A_X_B_* and males carrying one of three karyotypes each lacking a coherent *X_B_* segment.

### Recessive lethal structural variation segregating in Bunonematidae sp. JU3390

In Bunonematidae sp. JU3390 we identified a 988kb segregating recessive lethal segment on Scaffold 5 (Figure 5A, Supplementary Figure 4). Individuals carrying this segment lacked orthologous genes that in *C. elegans* are essential for normal development including genes orthologous to Structural Maintenance of Chromosomes (*smc*) isoforms *b* and *c* ^35^ and the highly conserved Jumonji Domain-containing Protein 2 *jmjd-2* ^36^ (Figure 5B). JMJD-2 demethylates lysine residues including H3K9 and H3K36, playing a critical role in the regulation of gene expression. Analysis of males and females identified no individuals homozygous for this deletion (Supplementary Figure 5). Instead, individuals carried multiple variants in this region including three that were separately assembled in our haplotype genome sequences (Supplementary Figure 4). Alignment of Scaffolds 2, 3, 4 and 6 did not identify other segregating variants of this large size (Supplementary Figure 6).

**Figure 5:**
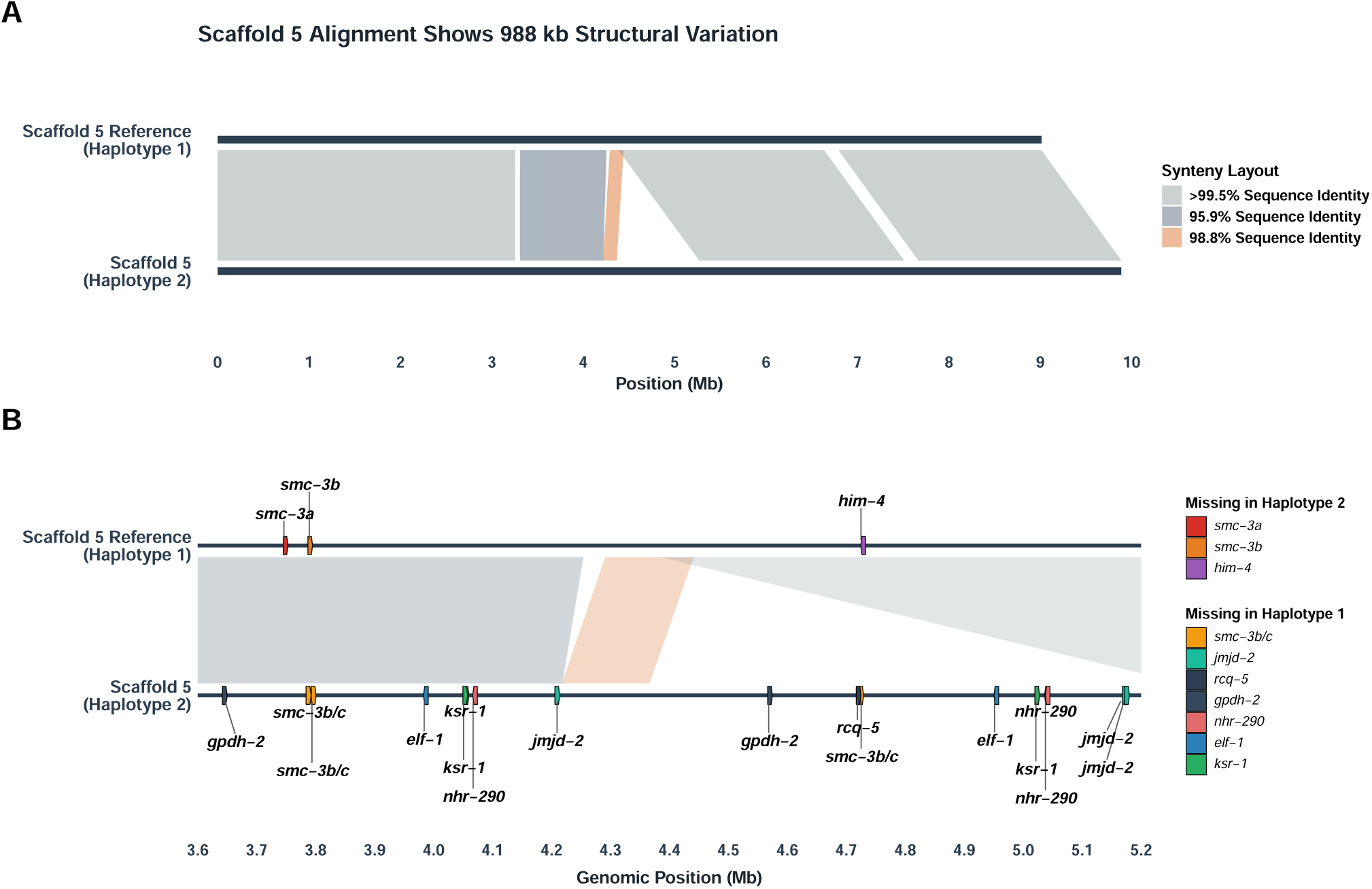
In Bunonematidae sp. JU3390 the Haplotype 1 and Haplotype 2 variants of Scaffold 5 showed A) significant structural variation with a 988 kb portion of Haplotype 2 not present in Haplotype 1. Analysis of protein-coding gene annotations in these region identified multiple genes with *C. elegans* orthologous partners that were present in Haplotype 2 but missing in Haplotype 1.

### Sex determination components downstream of TRA-1 are conserved in Bunonematidae sp. JU3390

The *X_B_* segment we identified in female Bunonematidae sp. JU3390 is large and contains thousands of annotated protein-coding sequences. In order to investigate the molecular basis of Bunonematidae sp. JU3390 sex determination we searched for components of the *C. elegans* sex determination network. Genes involved in sex determination in *C. elegans* have been extensively studied and many of the regulatory interactions are well-understood^37–39^. A number of individual genes coordinate molecular information to ‘count’ the X to autosome ratio, promote spermatogenesis or oogenesis, and participate in germline and somatic sex determination and differentiation. For example, the gene *tra-1* plays a critical role in sexual differentiation, actively repressing and regulating genes that promote male fate in *C. elegans* ^40^. Mutating *tra-1* causes XX individuals to develop as males instead of hermaphrodites.

We found molecular components upstream of *tra-1* are poorly conserved in both Bunonematidae sp. JU3390 and JU3391 (Figure 6A; Supplementary Tables 3-4). Most of the *C. elegans* proteins lacked orthologous counterparts in Bunonematidae sp. JU3390 or resulted in only weak potential alignments. Collectively, the regulatory interactions that constitute sex determination upstream of *tra-1* are not possible. In contrast, *tra-1* itself and downstream components were largely conserved in Bunonematidae sp. JU3390. The *C. elegans* TRA-1 protein includes five C2H2 zinc finger DNA binding motifs and a C-terminus that is cleaved and used in regulatory interactions^40^. Alignment of the *C. elegans* TRA-1 protein with Bunonematidae sp. genomic sequences identified an N-terminus region and high conservation of four of the five zinc finger DNA binding motifs. Both DNA alignment and annotated gene models suggested a truncated form of the protein in Bunonematidae sp. JU3390, lacking the C-terminus. Analyses in Bunonematidae sp. JU3391 identified similar TRA-1 molecular conservation but few conserved TRA-1 targets or germline effectors (Supplementary Table 4). Losing sexual reproduction eliminates the need for networks involved in male fates and mutational degradation of these components likely reflects the Bunonematidae sp. JU3391 transition to asexuality.

**Figure 6:**
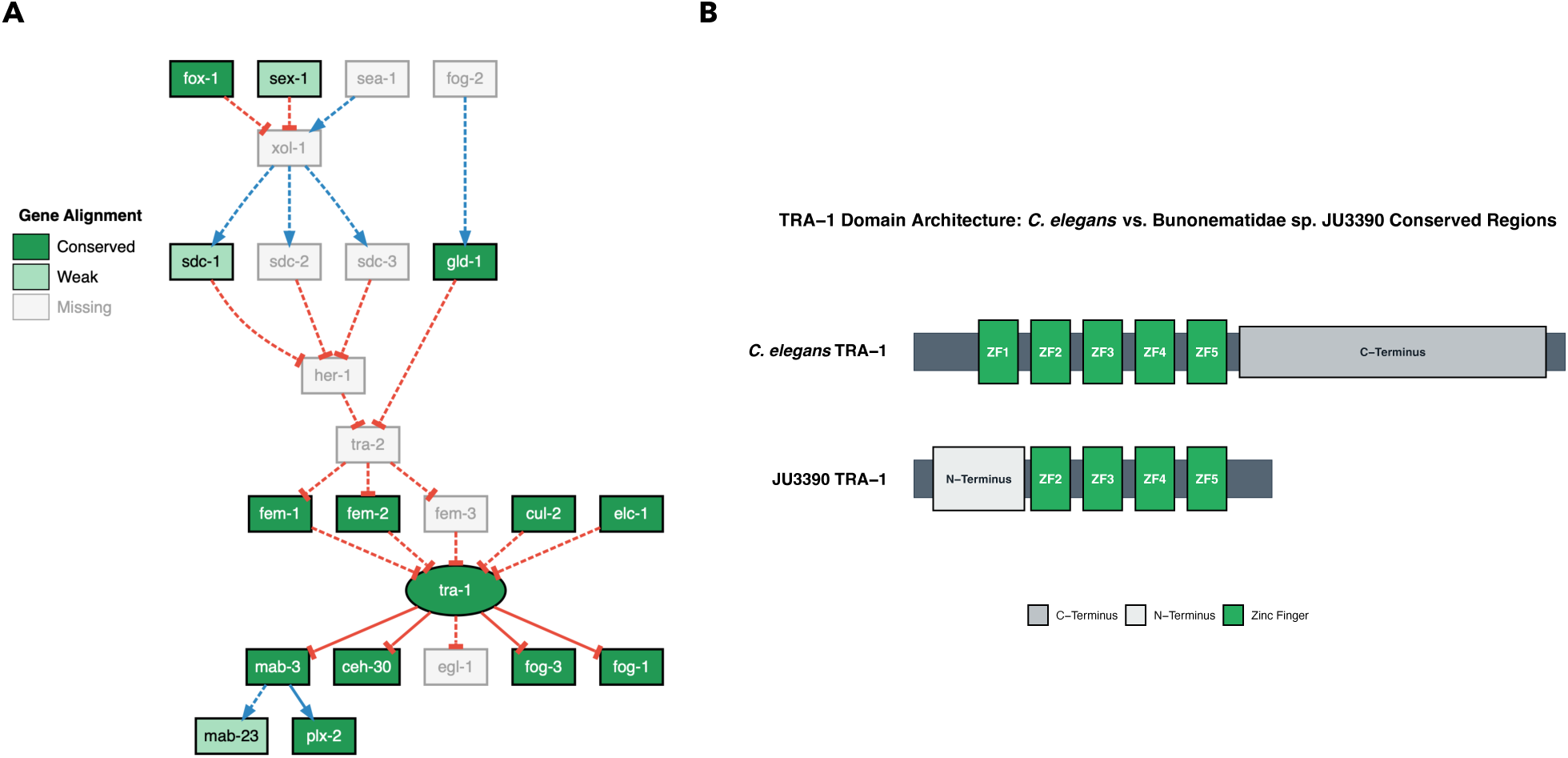
Orthologous genes in the *C. elegans* sex determination pathway that are upstream of TRA-1 are poorly conserved in Bunonematidae sp. JU3390. Orthologous TRA-1 and downstream components are well-conserved. A) Analysis of annotated protein-coding sequences and tblastx alignments between *C. elegans* genes involved in sex determination and Bunonematidae sp. JU3390 resulted in few conserved genes. The regulatory interactions present in *C. elegans* sex determination are not possible in Bunonematidae sp. JU3390 due to missing genes and broken regulatory relationships. In contrast, an orthologous TRA-1 and its downstream partners are conserved in Bunonematidae sp. JU3390. B) Bunonematidae sp. JU3390 TRA-1 and *C. elegans* TRA-1 share four of five zinc finger DNA binding motifs.

Of the conserved orthologous sex determination proteins, TRA-1, FOG-3 and FEM-2 were found on one of the Bunonematidae sp. JU3390 X variants. Both TRA-1 and FEM-2 were found on Scaffold 1 in Bunonematidae sp. JU3391 but there was only a weak potential FOG-3 alignment on Scaffold 5, suggesting this orthologous protein was not conserved. In *C. elegans* FOG-3 is repressed by TRA-1^41^ and promotes spermatogenesis in both hermaphrodites and males. Mutating FOG-3 in either sex transitions germ cells from sperm to oocytes^42^. FEM-2^43^ coordinately regulates TRA-1 with FEM-1 and FEM-3^44^, binding TRA-1 for degradation through the CUL-2/ELC-1 ubiquitin ligase pathway^45^. This degradation prevents TRA-1 from repressing male-specific factors including MAB-3 and FOG-3.

Orthologs of both TRA-1 and its downstream partner FOG-3 were found on Scaffold 1, *X_A_* and *X_B_* while competent FEM-2 orthologs were found only on Scaffold 1 and *X_B_* (Supplementary Table 3). Males carrying Scaffold 1 and *X_A_* have two copies of TRA-1 and FOG-3 and one copy of FEM-2 while females have three copies of TRA-1 and FOG-3 and two copies of FEM-2. In *C. elegans* FEM-2 promotes male specific fates and spermatogenesis but the Protein Phosphatase 2C family it belongs to is conserved across humans, yeast and *Arabidopsis* with no role in sex determination in these groups^43^. Thus, we find multiple potential causes of sexual fate in Bunonematidae sp. JU3390. Female fates may be governed by quantitative relationships between copy number variants, differential gene regulation or other forms of dosage balance. Alternately, these orthologous genes may have divergent roles in sex determination from those studied in *C. elegans* and other well-characterized nematodes, utilizing instead novel genes, interactions and regulatory mechanisms.

### Contingency and the re-invention of chromosomal sex

We hypothesize these volatile genome dynamics and novel sex chromosome system resulted from Bunonematidae sp. JU3390 worms escaping a build-up of lethal structural mutations. Evolutionary theory has struggled with the long-term persistence of asexual reproduction, suggesting that irreversible, eventually fatal ‘mutational meltdown’ in these species is inevitable^22,24^. Sexual outcrossing and the maintenance of variation and heterozygosity are a rapid means of escaping segregating deleterious variants that, if homozygous, would result in lethality. In Bunonematidae sp. JU3390 multiple large structural mutations such as those segregating on Scaffold 5 may have created a situation where population survival depended on finding a path out of parthenogenesis.

A fundamental question in biology is the role of contingency in evolution^46^. How much of the biodiversity that surrounds us is demanded by concrete, deterministic biology, and how much has been determined by contingency and chance^47^? Here we find the evolutionary history of Bunonematidae sp. nematodes points to a dominant role for contingency underpinning biodiversity. An evolutionary history of large structural mutations and potentially lethal homozygosity narrowed the worm’s paths for survival. In escaping this structural mutation ratchet Bunonematidae sp. JU3390 worms did not re-evolve sex; they re-invented it. The ancestral sex chromosome retired and a new chromosome underwent sex capture. The species did not return to the conserved nematode XY system, instead evolving a novel ZW sex chromosome system. Genome sequencing has provided an incredible lens into biology. The evolution of Bunonematidae sp. JU3390 nematodes and its re-invented sex system suggest we have only just started to discover the true diversity of life.

### Limitations of the study

Our bioinformatic and comparative genomic approach provides a high-resolution analysis of Bunonematidae sp. JU3390 and JU3391 but our study has several limitations. Assigning sex without destroying the animal is challenging in these microscopic worms and this, combined with the potential for environmental or other inputs to sex determination observed in related nematodes, means we can not conclusively identify the relationship between genomic content and sex in Bunonematidae sp. JU3390. Phylogenetic reconstruction and orthologous gene analysis suggest a history of parthenogenesis but as with all evolutionary reconstruction we can not conclusively verify this. Our PacBio and HiC data rely on sequencing pooled populations of thousands of worms and we can not yet reconstitute individual genomic content at high resolution. Our chromatin contact maps strongly suggest fusion of *X_A_* and *X_B_* segments but we can not verify this without cytology and experimental karyotyping. Similarly, our analysis of the components of the *C. elegans* sex determination network in Bunonematidae sp. JU3390 assumes conserved functions but requires experimental validation and characterization for conclusive results. Throughout, we have attempted to mitigate these limitations with high-quality, high-depth DNA sequence libraries, rigorous bioinformatic analyses and multiple approaches to validating results. Our bioinformatic and comparative genomic approach results in exciting possibilities for future experimental studies.

## RESOURCE AVAILABILITY

### Lead contact

Requests for further information and resources should be directed to and will be fulfilled by the lead contact, Janna L. Fierst.

### Materials availability

Nematode strains are available from the laboratory of Marie-Anne Felix through https://justbio.com/tools/worms-db/.

### Data and code availability

DNA and RNA data associated with this manuscript have been deposited with the National Center for Biotechnology Information under Bioproject PRJNA1501602 (JU3390 SAMN61944316; JU3391 SAMN61944317).

Scripts, bioinformatic workflows and documented analyses are available at https://github.com/jannafierst/HiC Assemblies.

The Arima genomics script is available at https://github.com/ArimaGenomics/Scripts/blob/master/generate site positions Arima.py.

## ACKNOWLEDGMENTS

The authors gratefully acknowledge Marie-Anne Felix and Aurelien Richaud for providing nematode strains, help and advice. This research was supported by NIGMS award R35147245 to JLF.

## AUTHOR CONTRIBUTIONS

JLF conceived the study and performed comparative genomic analyses. LAH, CAB, MAM and JAS cultured the nematodes, performed laboratory experiments, and extracted DNA and RNA for sequencing. PK performed single worm lysis and MDA. VKE and KKW performed bioinformatic analyses. Each of the authors contributed text, edits and feedback for the manuscript.

## DECLARATION OF INTERESTS

The authors declare no competing interests

## DECLARATION OF GENERATIVE AI AND AI-ASSISTED TECHNOLOGIES

During the preparation of this work, the authors used Gemini AI to produce generalized, commented UNIX scripts from rough drafts. The authors also used Gemini AI to produce rough draft R scripts for plotting, visualization and latex table formatting. After using Gemini AI for these services, the authors reviewed and edited the content as needed and take full responsibility for the content of the publication.

## SUPPLEMENTAL INFORMATION INDEX

Figures S1-S7 and their legends in a PDF

Table S1. Bunonematidae sp. JU3390 and JU3391 sequence library statistics

Table S2. Bunonematidae sp. JU3390 and JU3391 sequence assembly statistics

Table S3. Genes in the sex determination pathway in *C. elegans* and their orthologous counterparts in Bunonematidae sp. JU3390

Table S4. Genes in the sex determination pathway in *C. elegans* and their orthologous counterparts in Bunonematidae sp. JU3391

## STAR METHODS

### Experimental model and study participant details

#### Strains and culturing

We obtained Bunonematidae sp. strains JU3390 and JU3391 from the laboratory of Marie-Anne Felix in Paris, France. The worms were collected from rotting grass, leaves and leaf litter near the University of Galway, Ireland on July 1, 2018 (geographic details are given on justbio.com/tools/worms-db/). Worms were maintained on standard Nematode Growth Medium (NGM) plates with added Nystatin and Streptomycin, seeded with *Escherichia coli* OP50-1 at 20*^◦^*C^48^.

### Method details

#### DNA and RNA extraction and sequencing

We extracted high molecular weight (HMW) DNA from pooled populations of mixed stage worms. Both strains were grown on three 100mm NGM plates each for two weeks, which was sufficient time for the nematodes to fill the plates. Worms were washed off plates with 1x M9 buffer into 15 mL conical tubes and washed twice with 1x M9 buffer to remove superficial OP50-1. Worms were then left on a rocker overnight (approximately 17 hours) to mitigate the amount of OP50-1 in their guts, then washed twice more with 1x M9. Worms were then pelleted by centrifugation and after decanting, the pellet was divided into 50 *µ*L aliquots in 1.5 mL tubes. One aliquot per strain was used for HMW DNA extraction. Prior to extraction, nematode cuticles were broken with repeated freeze-thaw cycles, where the tubes were placed at -80*^◦^*C for five minutes, moved to 37C until thawed, then moved back to -80*^◦^*C for a total of five cycles. We then used the Promega Wizard HMW DNA Extraction Kit (Promega, Cat. #A2920), adapting the manufacturer’s suggested protocol for HMW DNA isolation from tissue culture cells. Specifically, working volumes of isopropanol and 70% ethanol were kept on ice throughout the protocol, all centrifugation steps were performed at 4*^◦^*C and 16,000 *x g*, and the cell lysis step was augmented with a 25-minute incubation at 65*^◦^*C followed by 1 minute on ice.

Each DNA library was sequenced on a Pacific Biosystems Sequel II instrument at the University of Miami Center for Genome Technology to generate highly accurate HiFi reads^49^. HiC DNA libraries were produced by Arima Genomics in San Diego, California from one 50 *µ*L aliquot of frozen mixed stage worms and sequenced on an Illumina NovaSeq 6000^30^.

Culturing for RNA extraction was performed exactly as for DNA extractions, and nematodes were washed using the same methods. After pelleting and decanting, 10 *µ*L of live nematode pellet was flash-frozen on liquid nitrogen. Tissues were disrupted using the Cellcrusher-mini system (https://cellcrusher.com/mini-tissue-pulverizer-3/). We used the QIAGEN RNeasy Mini Kit (QIAGEN, Cat. #74104) following manufacturer protocol. Samples were sent on dry ice to GENEWIZ (Azenta Life Sciences) and sequenced on an Illumina NovaSeq 6000^30^.

#### Single-worm lysis, multiple displacement amplification, and Illumina sequencing

Whole-genome amplification from one intact adult worm per sample was performed through phi29 DNA polymerase-based multiple displacement amplification (MDA)^50^ using the REPLI-g Mini Kit (QIAGEN, Cat. #150023). The procedure was adapted from the single-worm MDA method of Lee et al.^51,52^ and the manufacturer’s protocol^53^. Prior to lysis, worms were washed as described for whole population DNA and RNA extractions. The sex of each individual was called phenotypically by presence/absence of a spicule, a male mating structure visible both when extended and retracted. Because retracted spicules are difficult to see, these calls have some margin of error. Each worm was processed whole without cutting or puncturing.

Buffer DLB was reconstituted with 500 *µ*L of nuclease-free water. Fresh Buffer D2 was prepared by mixing 1 M dithiothreitol with reconstituted Buffer DLB at a 1:11 ratio, v/v. A single intact worm was transferred into a 0.2 mL PCR tube containing 4 *µ*L of 1*×* PBS. Thereafter, 3 *µ*L of Buffer D2 was added and the sample was incubated for 10 minutes at room temperature, followed by 10 minutes at 65*^◦^*C with the heated lid maintained at no more than 70*^◦^*C. Then 3 *µ*L of REPLI-g Stop Solution was added and placed the resulting 10 *µ*L crude lysate on ice.

For MDA, a 40 *µ*L master mix containing 10 *µ*L of nuclease-free water, 29 *µ*L of REPLI-g Mini Reaction Buffer, and 1 *µ*L of REPLI-g Mini DNA Polymerase was prepared. The master mix was added to the entire 10 *µ*L crude lysate, producing a final reaction volume of 50 *µ*L. MDA proceeded at 30*^◦^*C for 16 hours. The DNA polymerase was subsequently inactivated at 65*^◦^*C for 3 minutes with the heated lid set to 70*^◦^*C. Amplified DNA was stored at -20*^◦^*C and quantified using the Qubit dsDNA High-Sensitivity Assay Kit (Invitrogen, Cat. #Q32854) according to the manufacturer’s instructions.

Amplified DNA was purified using AMPure XP paramagnetic beads (Beckman Coulter Life Sciences, Cat #A63882). 90 *µ*L of resuspended beads were added to 45-50 *µ*L of amplified DNA and the samples were mixed on Hula Mixer for 5 minutes at room temperature. The tubes were placed on a magnetic rack, and the supernatant was removed after the beads collected against the tube wall. The beads were washed two to three times with 500 *µ*L of freshly prepared 80% ethanol. After the final wash, residual ethanol was removed, and the beads were air-dried without allowing the pellet to crack. DNA was eluted in 50 *µ*L of nuclease-free water at 37°C for 35 minutes. Following magnetic separation, the clear eluate was transferred to a new tube without disturbing the beads. Purified DNA was quantified using the Qubit dsDNA High-Sensitivity Assay Kit. Purified MDA products were submitted to GENEWIZ for whole genome Illumina sequencing.

#### Genome assembly, decontamination and scaffolding

We assembled genome sequences from Pacific Biosystems Hifi and Arima HiC libraries with the Hifiasm assembly software v0.25^54^. We initially ran hifiasm with phased diploid assembly for both JU3390 and JU3391 (for example; `hifiasm -o JU3390 -t 32 --h1 JU3390 arima R1.fastq --h JU3390 arima R2.fastq JU3390 hifi.fastq`) where -o is the output directory, -t specifies threads --h1 and --h2 specify arima HiC libraries and JU3390 hifi.fastq is the Pacific Biosystems Hifi DNA library. Alignment of our Hifi library to each of the two phased JU3390 haplotypes with Minimap2 v2.30-r1287^55^ resulted in low MAPQ scores for Scaffold 1 and we also assembled JU3390 triploid and tetraploid haplotypes with `--n-hap 3` and `--n-hap 4`. The tetraploid assembly produced two differentiated haplotypes and two haplotypes identical in number and length of contiguous sequences and we selected the triploid assembled haplotypes for reference sequence analysis.

Nematode cultures contain significant microbial contamination and part of the sequencing library and assembled contiguous sequences were contaminants of microbial origin. To identify these we used BLAST-PLUS v.2.16.0^56^ to align sequences to the NCBI nucleotide (nt) database (v5; downloaded March 10, 2025). Contiguous sequences were randomly sampled for 10kb regions to accelerate identification and if no alignments were identified we re-attempted BLAST alignment for that contiguous sequence. Contiguous sequences with identified nematode origins or unidentified origins were retained in the assembled genome sequence and those with identified microbial origins were discarded.

We used the purge dups software v1.2.6^57^ to eliminate allelic regions from the assembled genome sequence. Briefly, we aligned the Hifi reads to the assembled sequences with Minimap2 v2.30-r1287^55^ to estimate sequence read coverage, used the purge dups pbcstat module and calcuts to estimate sequencing coverage, self-aligned the assembled sequences with Minimap2 v2.30-r1287^55^ to identify duplications and used these estimates to purge duplicated and allelic regions (for example; `purge dups -2 -T cutoffs -c PB.base.cov JU3390.split.self.paf.gz > dups.bed 2>purge dups.log`). We extracted sequences with the get seqs module and retained the purged haplotigs as the assembled, decontaminated, purged haplotype.

We used the BUSCO software v5.4.3^58^ and the nematoda odb12 lineage dataset to analyze completeness of each genome sequence before and after allelic removal (Extended Data Table 2). We used trim galore v0.6.10^59^ with the options `--paired --quality 20 –stringency 3 --length 20 --fastqc --cores 32` to remove internal adapters and discard low-quality sequences in the HiC libraries. We used the Arima genomics generate site positions Arima.py python script with the cut sites options `-e GATC GANTC TTAA CTNAG` to identify potential enzymatic cut sites in the assembled genome sequences. We then aligned the HiC libraries to the contiguous sequences with the bwa alignment software v0.7.19-r1273^60^ within the Juicer v2.17.00 software^61^. The resulting decontaminated, purged sequences were further assembled into scaffolds with the YaHS software v1.2.2^62^ and default configuration (for example; `yahs JU3390 haplotype 1 JU3390 haplotype 1.bam`) where the .bam file is the output of the Juicer / bwa alignment pipeline.

### Quantification and statistical analysis

#### Phylogenetic reconstruction

We reconstructed the Rhabditina phylogeny following the methods described in^63^. Briefly, we extracted 70 universal single-copy orthologs from the assembled genome sequences of 101 Rhabditina nematode genomes, aligned with MAFFT^64^, trimmed with ClipKIT^65^ and estimated phylogenetic distances and relationships with IQTREE^66^. Visualization and plotting were done in RStudio 2026.06.0+242 with the software ape^67^, phytools^68^ and ggplot2 with the R software version 4.5.3 ”Reassured Reassurer.”

#### Annotation

We annotated protein-coding genes with the BRAKER4 software^58,69–82^. Prior to BRAKER4, we aligned mRNA libraries to the assembled scaffolds with the STAR software^83^ with option – outSAMstrandField intronMotif, and we created repeat-masked genome sequences with the RepeatMasker^84^ and RepeatModeler2 software^85^. The BAM from STAR and softmasked genome from RepeatMasker were input to BRAKER4, which was run with default parameters with exception of option downsampling lambda = 1 for JU3390, and downsampling lambda = 0 for JU3391. Smaller lambda values more aggressively downsample single-exon genes during training. Variations of lambda were tested and these gave the most complete gene sets according to Compleasm^72^ and BUSCO^58^. Longest isoforms in our predicted protein-coding annotations were analyzed for functional regions using InterProScan software^86^. Longest isoforms were also input to OrthoFinder^87^ in a pairwise fashion between JU3390, JU3391, and the WormBase *C. elegans* genome published in 2014. Single-copy orthologs from OrthoFinder were used to investigate Nigon elements and chromosome identity of the scaffolds following^16^.

### Alignment and analysis

We used the Minimap2 software^55^ to align Pacific Biosystems Hifi libraries to the scaffolded chromosomes with `-ax map-hifi` and used DeepVariant v1.6.0^88^ to identify segregating genomic variants with `MODEL TYPE "PACBIO"`. To assign karyotype identity we combined aligned Hifi sequencing coverage across each genomic segment with sequencing coverage across breakpoints identified in Scaffold 1, *X_A_* and *X_B_*. We first aligned individual MDA Illumina DNA libraries to each of the assembled haplotypes with minimap2, excluding secondary alignments -`ax sr -N 1 --secondary=no` and estimated sequencing coverage with samtools depth `-a -Q 0 -G SECONDARY`. We measured mapping quality through MAPQ scores where high values indicate high confidence in alignment.

We extracted sequence reads aligned to the breakpoints identified in Scaffold 1, *X_A_* and *X_B_* alignment (Fig. 2E) and measured the total number of sequence reads aligned to the region, the number that were aligned continuously without soft clipping. For Haplotype 1 these were Scaffold 1 1708900-1709100 and 3650900-365110; for Haplotype 2 these were *X_A_* 7929597-7929797 and *X_B_* 3693563-3693763 and for Haplotype 3 these were *X_A_* 7923700-7923900 and *X_B_* 3693563-3693763. We then assigned a concordance score for each genomic segment where Concordance = (100 - percent of aligned reads with a MAPQ of 0) * (percent of aligned reads continuously covering the breakpoint * 100). Here, MAPQ 0 indicates a sequence read maps equally well to two or more regions in the haplotype.

### Identification of genes orthologous to those in the *C. elegans* sex determination network

We curated a list of genes and regulatory interactions in the *C. elegans* sex determination network from WormBook^37,38^ (Supplementary Table 3). We obtained protein-coding sequences for each of these genes from NCBI and used TBLASTN^56^ to search for potential sequences in the Bunonematidae sp. JU3390 and JU3391 assembled chromosomes. We validated putative alignments with OrthoFinder results^87^, searching orthogroups for *C elegans* sex determination genes. Each orthologous gene was marked ‘Present’ if we identified an orthogroup containing the *C. elegans* protein and a Bunonematidae sp. JU3390 or JU3391 annotated protein-coding gene and a TBLASTN^56^ alignment in the same region of the genome with bitscore *>*50. Genes were marked ‘Weak’ if there were no annotated protein-coding genes and a TBLASTN^56^ alignment with bitscore *>*50 and ‘Missing’ if there were no annotated protein-coding genes and the TBLASTN^56^ alignment had bitscore *<*50. To identify conserved regions of the TRA-1 protein we aligned the *C. elegans* TRA-1 sequence to the Bunonematidae sp. JU3390 and JU3391 assembled sequences using exonerate with a protein2genome model^89^.

## Notes

https://github.com/jannafierst/HiC_Assemblies

### Competing Interest Statement

The authors have declared no competing interest.

### Summary of Updates

Added new analyses, corrected species name references throughout

