## Supplementary Figures and Tables for "Re-inventing sex: A W chromosome and a path out of parthenogenesis for Bunonematidae sp. nematodes"

#### SUMMARY

Supplementary Figures

#### KEYWORDS

evolution, genetics, sex chromosomes, nematodes, contingency

#### Supplementary Figures

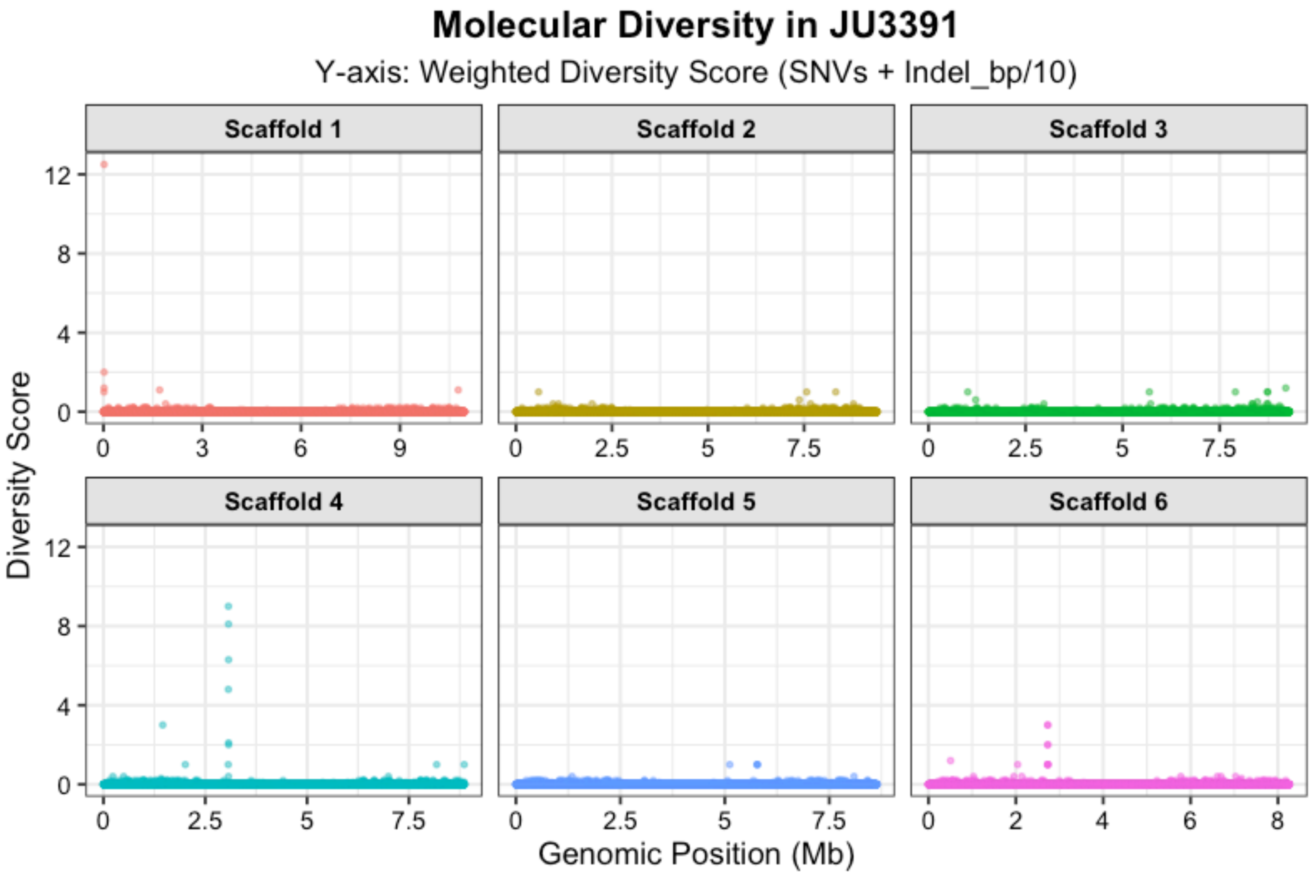


Figure 1: Molecular diversity in Bunonematidae sp. JU3391 is extremely low with just 1,738 SNVs across the entire genome and an effective heterozygosity rate of 0.003%.

#### Single Copy Orthologous Genes and *C. elegans* Chromosome Locations

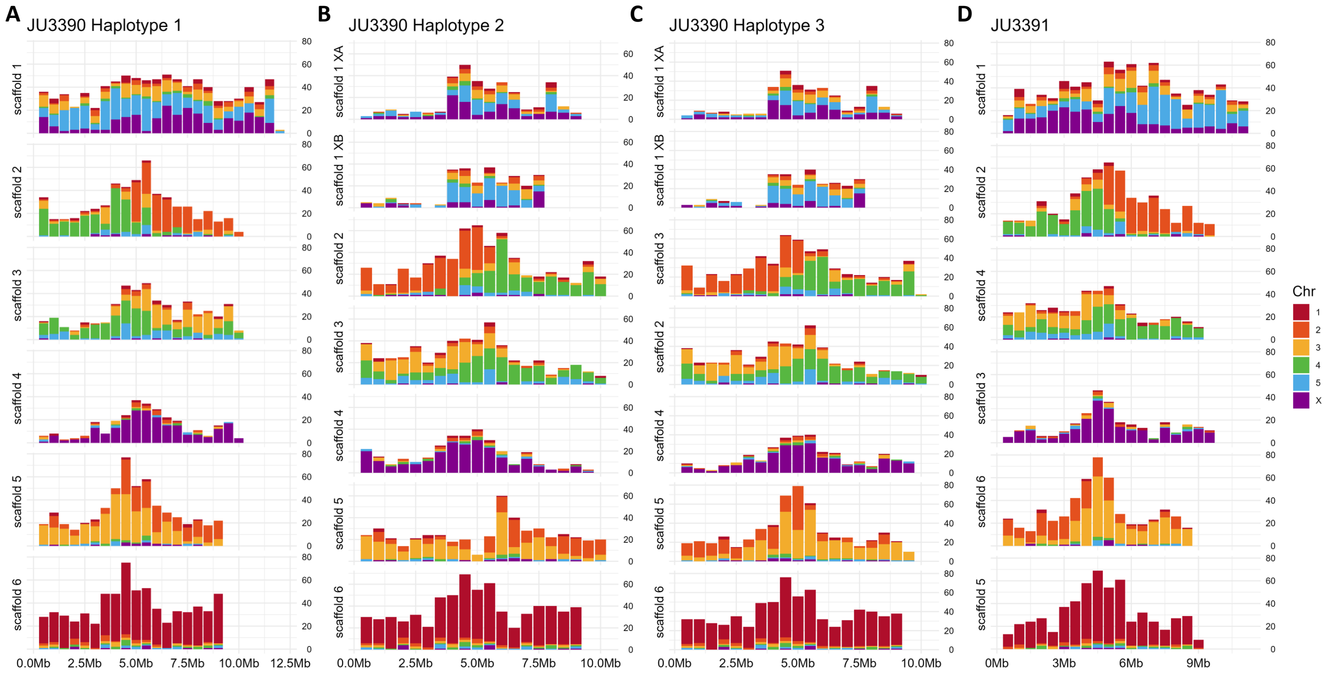


Figure 2: Single copy orthologous genes across A-C) Bunonematidae sp. JU3390 and D) JU3391 showed lack of synteny with *C. elegans* chromosomal locations. The bulk of genes found on the X chromosome in *C. elegans* were found on Scaffold 1, $X_A$, $X_B$ and Scaffold 4 in Bunonematidae sp. JU3390 and Scaffold 1 and Scaffold 3 in Bunonematidae sp. JU3391.

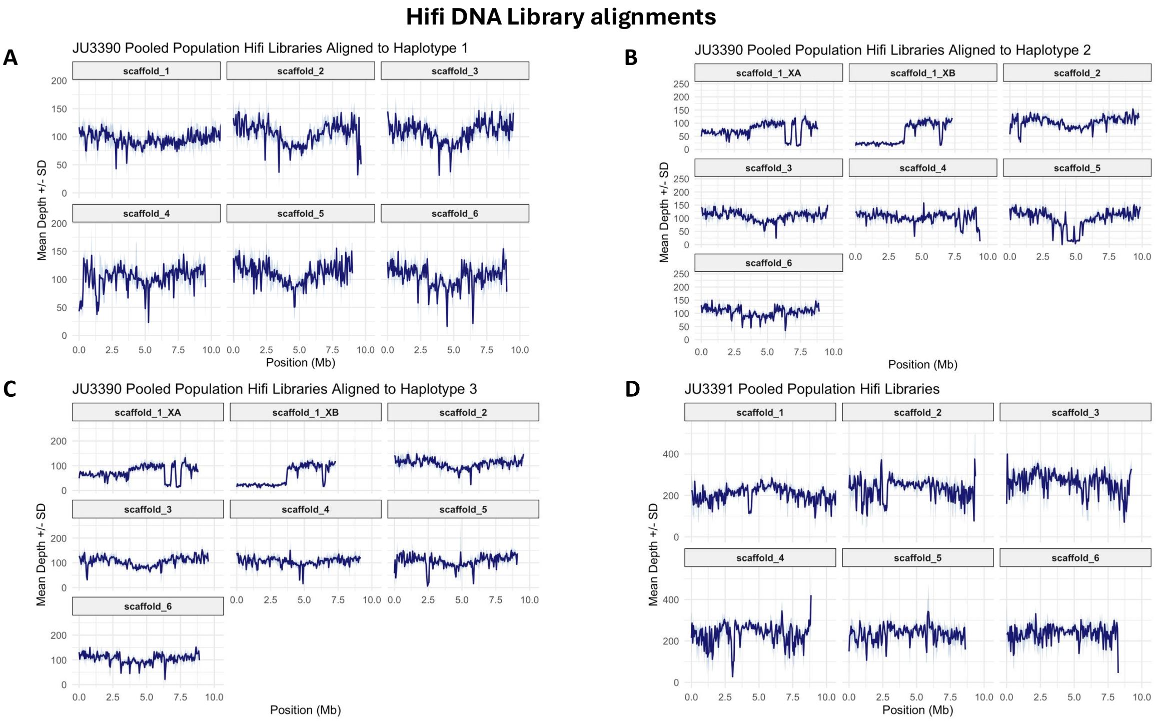


Figure 3: Alignment of Hifi DNA sequence libraries to Bunonematidae sp. JU3390 Haplotype 1 showed significant variance across the assembled scaffolds. In B) Haplotype 2 and C) Haplotype 3 high homology between the $X_A$ and $X_B$ segments prevented unique alignment. In Bunonematidae sp. JU3391 alignment of Hifi DNA sequence libraries to the assembled scaffolds demonstrated similar variance in coverage.

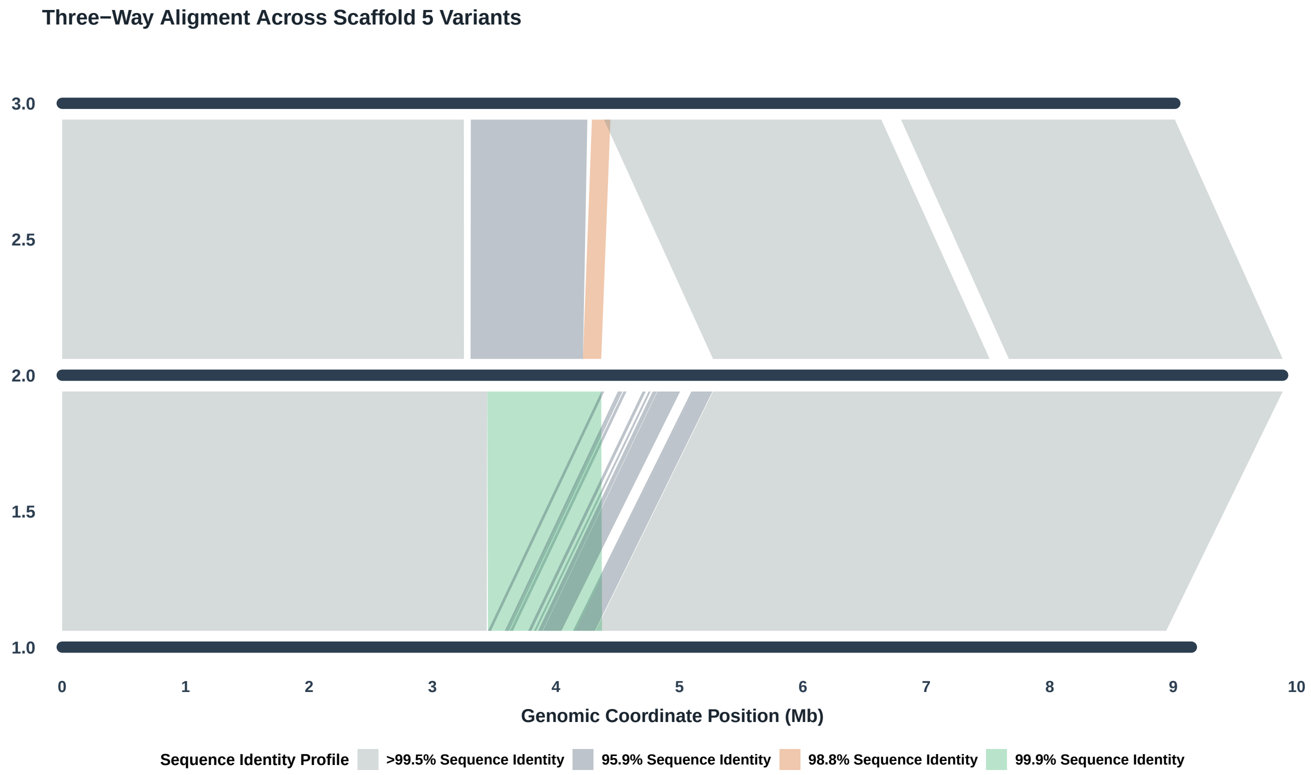


Figure 4: Alignment of Scaffold 5 across the three Bunonematidae sp. JU3390 haplotypes demonstrated that the 988kb unique region present in Haplotype 2 aligned to multiple tandem variants in Haplotype 3.

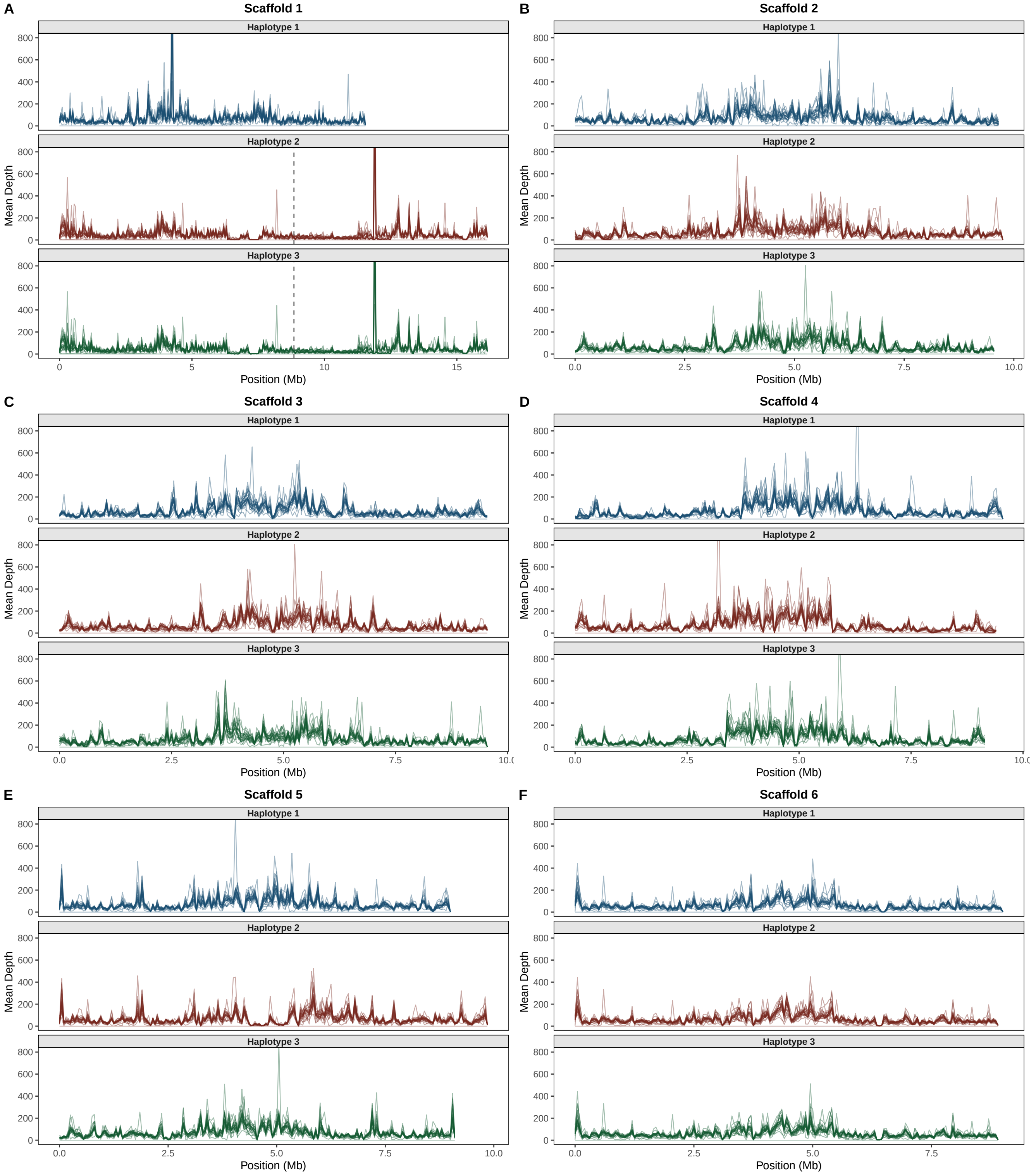


Figure 5: Single worm IIllumina DNA libraries generated with MDA aligned across A-F) the six scaffolds of haplotypes 1, 2 and 3. The alignments indicate the libraries successfully captured the assembled genome sequence but coverage was extremely noisy and variable across all assembled sequences.

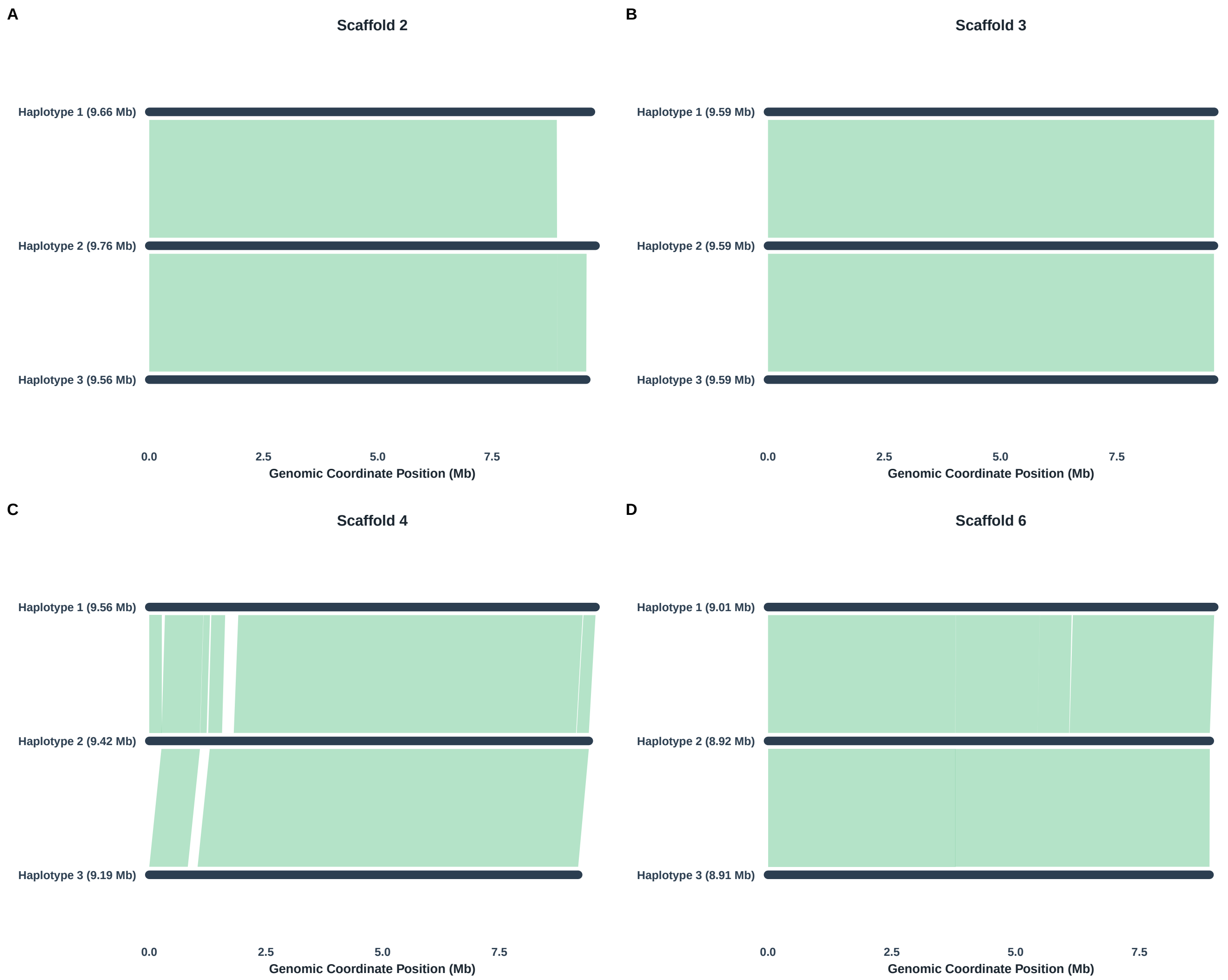


Figure 6: Alignment of A-D) Scaffolds 2, 3, 4 and 6 across the 3 assembled haplotypes did not identify large segregating variants like those in Scaffolds 1 and 5

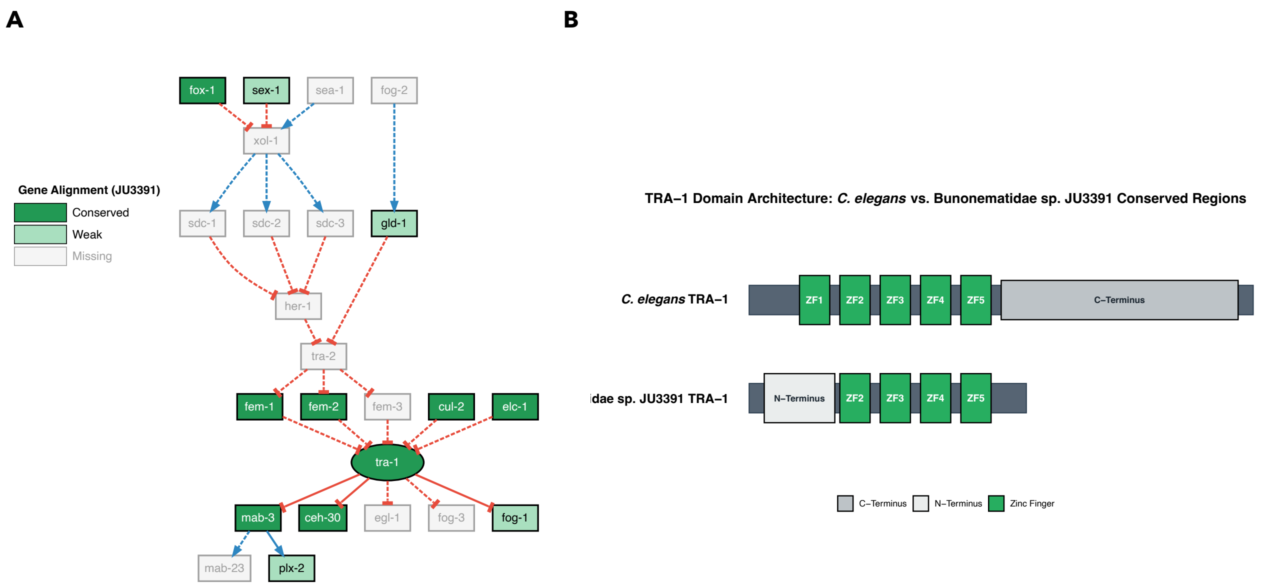


Figure 7: Bunonematidae sp. JU3391 orthologous genes in the *C. elegans* sex determination pathway are less well-conserved when compared with Bunonematidae sp. JU3391

**Table 1. JU3390 and JU3391 Sequence Library Statistics**

| | JU3390 | JU3391 |
|---|---|---|
| Hifi Data (Library N50) | 20.5Gb (15.3Mb) | 19.2Gb (15.5Mb) |
| HiC Data | 54.0Gb | 44.0Gb |
| mRNA-Seq data | 3.9Gb | 6.9Gb |

#### Table 2. Genome Assembly Statistics

| | | JU3390 Haplotype 1 | JU3390 Haplotype 2 | JU3390 Haplotype 3 | JU3391 p |
|---|---|---|---|---|---|
| BUSCO (genome) | Hifiasm assembly | C:92.1%[S:68.0%,D:24.2%], F:1.0%,M:6.9%,n:596 | C:92.1%[S:65.1%,D:27.0%], F:1.2%,M:6.7%,n:596 | C:92.1%[S:65.1%,D:27.0%], F:1.2%,M:6.7%,n:596 | C:93.0%[S:75.3%,D:17.6%], F:0.8%,M:6.2%,n:596 |
| | Decontaminated assembly | C:92.1%[S:68.0%,D:24.2%], F:1.2%,M:6.7%,n:596 | C:92.1%[S:64.9%,D:27.2%], F:1.2%,M:6.7%,n:596 | C:92.1%[S:64.9%,D:27.2%], F:1.2%,M:6.7%,n:596 | C:92.8%[S:75.7%,D:17.1%], F:1.2%,M:6.0%,n:596 |
| | Purged of haplotigs assembly | C:92.1%[S:91.6%,D:0.5%], F:1.2%,M:6.7%,n:596 | C:92.1%[S:75.5%,D:16.6%], F:1.2%,M:6.7%,n:596 | C:92.1%[S:75.5%,D:16.6%], F:1.2%,M:6.7%,n:596 | C:92.8%[S:91.9%,D:0.8%], F:1.2%,M:6.0%,n:596 |
| | Scaffolded assembly | C:92.1%[S:91.6%,D:0.5%], F:1.2%,M:6.7%,n:596,E:39.5% | C:92.1%[S:75.5%,D:16.6%], F:1.2%,M:6.7%,n:596,E:39.2% | C:91.9%[S:78.2%,D:13.8%], F:1.2%,M:6.9%,n:596 | C:92.8%[S:91.9%,D:0.8%], F:1.2%,M:6.0%,n:596,E:43.4% |
| Genome size (Mb) | Hifiasm assembly | 129.1 | 122.1 | 184.5 | 128.4 |
| | Decontaminated assembly | 71.7 | 72.4 | 72.6 | 73.4 |
| | Purged of haplotigs assembly | 64.0 | 64.1 | 64.0 | 56.7 |
| | Scaffolded assembly | 58.4 | 64.1 | 63.1 | 56.7 |
| # scaffolds | Hifiasm assembly | 273 | 207 | 1907 | 560 |
| | Decontaminated assembly | 45 | 59 | 65 | 372 |
| | Purged of haplotigs assembly | 26 | 25 | 26 | 39 |
| | Scaffolded assembly | 7 | 10 | 13 | 27 |
| N50 (Mb) | Hifiasm assembly | 4.3 | 4.3 | 2.3 | 3.9 |
| | Decontaminated assembly | 4.9 | 4.5 | 4.5 | 4.5 |
| | Purged of haplotigs assembly | 4.9 | 4.9 | 4.9 | 6.3 |
| | Scaffolded assembly | 9.6 | 9.4 | 9.1 | 9.3 |
| N90 (Mb) | Hifiasm assembly | 0.03 | 0.04 | 0.03 | 0.06 |
| | Decontaminated assembly | 1.6 | 1.6 | 0.7 | 0.05 |
| | Purged of haplotigs assembly | 2.5 | 2.5 | 2.5 | 2.3 |
| | Scaffolded assembly | 9.0 | 7.3 | 7.3 | 8.3 |
| L50 | Hifiasm assembly | 11 | 11 | 20 | 11 |
| | Decontaminated assembly | 5 | 6 | 6 | 5 |
| | Purged of haplotigs assembly | 5 | 5 | 5 | 4 |
| | Scaffolded assembly | 3 | 4 | 4 | 3 |
| L90 | Hifiasm assembly | 43 | 38 | 1281 | 226 |
| | Decontaminated assembly | 14 | 15 | 16 | 151 |
| | Purged of haplotigs assembly | 12 | 12 | 12 | 10 |
| | Scaffolded assembly | 6 | 7 | 7 | 6 |
| GC % | Hifiasm assembly | 52.17 | 52.56 | 51.42 | 52.01 |
| | Decontaminated assembly | 50.06 | 50.05 | 50.06 | 47.34 |
| | Purged of haplotigs assembly | 50.02 | 50.02 | 50.02 | 47.32 |
| | Scaffolded assembly | 50.01 | 50.02 | 50.00 | 47.32 |
| # genes (braker, longest isoform) | Scaffolded assembly | 10424 | 11280 | 11252 | 13943 |
| BUSCO (protein set) | Scaffolded assembly | C:92.4%[S:91.4%,D:1.0%], F:3.5%,M:4.0%,n:596 | C:92.1%[S:76.5%,D:15.6%], F:3.4%,M:4.5%,n:596 | C:93.1%[S:80.0%,D:13.1%], F:2.9%,M:4.0%,n:596 | C:89.3%[S:87.4%,D:1.8%], F:5.5%,M:5.2%,n:596 |
| Compleasm | Scaffolded assembly | C:94.63%[S:48.83%,D:45.81%] F:2.35%,M:3.02%,n:596 | C:94.63%[S:42.28%,D:52.35%] F:2.01%,M:3.36%,n:596 | C:94.8%[S:43.96%,D:50.84%], F:2.01%,M:3.19%,n:596 | C:90.78%[S:56.04%,D:34.73%], F:5.03%,M:4.19%,n:596 |
| OMArk | Scaffolded assembly | C:74.28%[S:71.79%,D:2.49%], M:25.72%,n:3052 | C:73.62%[S:65.24%,D:8.39%], M:26.38%,n:3052 | C:74.28%[S:66.38%,D:7.90%], M:25.72%,n:3052 | C:78.64%[S:73.17%,D:5.47%], M:21.36%,n:3052 |
| % repeats (repeatmasker) | Scaffolded assembly | 12.39% | 12.90% | 12.36% | 19.51% |
| % repeats (earlgrey) | Scaffolded assembly | 13.71% | 13.39% | 13.42% | 19.35% |
| # SSCOs to *C.elegans* (orthofinder) | Scaffolded assembly | 3651 | 3378 | 3379 | 3304 |

### Bunonematidae sp. JU3390 - *C. elegans* Sex Determination Pathway Conservation

| Gene | Role | Location | E-value | Bit | OrthoGroup | *C. elegans* | Status |
|---|---|---|---|---|---|---|---|
| **Upstream X:A Counting Network** | | | | | | | |
| fox-1 | X-chromosome signal | Scaffold 4 | 5.79e-21 | 93 | OG0000803 | NP_001317738.1 | Present |
| sex-1 | X-chromosome signal | Scaffold 3 | 7.36e-21 | 96.1 | – | NP_001024662.1 | Weak |
| sea-1 | Autosomal signal | Scaffold 5 | 1.75e-04 | 44.1 | – | NP_494611.1 | Missing |
| xol-1 | X:A Counting | No Hits | – | – | – | NP_001024418.1 | Missing |
| sdc-1 | Dosage compensation | Scaffold 1 | 8.41e-04 | – | OG0002300 | NP_001360585.1 | Weak[1] |
| sdc-2 | Dosage compensation | No Hits | – | – | – | NP_509924.1 | Missing |
| sdc-3 | Dosage compensation | No Hits | – | – | – | NP_506703.1 | Missing |
| **Somatic Sex Determination** | | | | | | | |
| her-1 | Secreted masculinizer | No Hits | 0.70 | – | – | NP_001024310.1 | Missing |
| tra-2 | Membrane receptor | Scaffold 2 | 0.006 | 42.1 | – | NP_495426.1 | Missing |
| fem-3 | Intracellular signaling | No Hits | 1.30 | 32.3 | – | NP_001255365.1 | Missing |
| **Signaling & Adapters** | | | | | | | |
| fem-1 | Intracellular signaling | Scaffold 2 | 2.21e-14 | – | OG0002468 | NP_500824.1 | Present |
| fem-2 | Intracellular signaling | Scaffold 1 / XB | 5.11e-12 | – | OG0004093 | NP_497224.1 | Present[2] |
| cul-2 | Ligase Scaffold | Scaffold 1 | 2.70e-19 | 93.9 | OG0001245 | NP_001023008.1 | Present |
| elc-1 | Ligase Adaptor | Scaffold 1 | 2.21e-23 | 90.8 | OG0002362 | NP_001380023.1 | Present[3] |
| zyg-11 | Cell Cycle / Meiosis | Scaffold 3 | 1.72e-09 | 62.4 | – | NP_495677.1 | Weak |
| **Somatic Sex Determination Regulator** | | | | | | | |
| tra-1 | Regulation of Sex Determination | Scaffold 1 / XB / XA | 6.50e-47 | – | OG0001328 | NP_001022880.2 | Present |
| **Germline Effectors** | | | | | | | |
| fog-2 | Germline switch | No Hits | 3.60 | – | – | NP_001041187.1 | Missing |
| gld-1 | Germline repressor | Scaffold 6 | 4.43e-25 | – | OG0014323 | NP_492143.1 | Present |
| fbf-1/2 | Germline fate | Scaffold 3 | 1.33e-50 | – | OG0000402 | NP_495216.1 | Present |
| fog-1 | Germline switch | Scaffold 3 | 3.75e-21 | – | – | NP_001367957.1 | Present |
| fog-3 | Germline switch | Scaffold 1 / XB / XA | 4.12e-05 | – | OG0004746 | NP_492646.1 | Present |
| **Somatic Male Targets** | | | | | | | |
| mab-3 | Somatic Male Switch | Scaffold 2 | 1.18e-25 | 72.2 | OG0002564 | NP_001022464.1 | Present |
| ceh-30 | Male Neuron Survival | Scaffold 2 | 6.88e-14 | 69.7 | OG0004232 | NP_508524.2 | Present |
| egl-1 | Apoptosis Activator | No Hits | – | – | – | NP_506575.2 | Missing |
| mab-23 | Male Ray Development | Scaffold 3 | 0.76 | 27.5 | OG0002354 | NP_001041090.1 | Weak |
| plx-2 | Axon Guidance | Scaffold 2 | 5.04e-16 | 74.7 | OG0000397 | NP_001364609.1 | Present |
| **Regulatory Partners** | | | | | | | |
| daf-16 | Aging / Diapause | Scaffold 6 | 1.81e-27 | 116 | – | NP_001367893.1 | Present |
| hen-1 | Nerve ring signaling | Scaffold 2 | 0.001 | 36.8 | – | NP_509200.1 | Missing |

[1]Weak hits and no gene models. [2]Copies on Scaffold 1 and XB but no XA. [3]Copy on Scaffold 1 only.

### Bunonematidae sp. JU3391 - *C. elegans* Sex Determination Pathway Conservation

| Gene | Role | Location | E-value | Bit | OrthoGroup | *C. elegans* | Status |
|---|---|---|---|---|---|---|---|
| **Upstream X:A Counting Network** | | | | | | | |
| fox-1 | X-chromosome signal | Scaffold 3 | 8.27e-20 | 90.8 | OG0001823 | NP_001317738.1 | Present |
| sex-1 | X-chromosome signal | Scaffold 4 | 1.14e-21 | 98.6 | – | NP_001024662.1 | Weak |
| sea-1 | Autosomal signal | Scaffold 6 | 0.002 | 40.7 | – | NP_494611.1 | Missing |
| xol-1 | X:A Counting | Scaffold 3 | 0.57 | 33.1 | – | NP_001024418.1 | Missing |
| sdc-1 | Dosage compensation | Scaffold 1 | 2.07e-04 | 46.6 | OG0002372 | NP_001360585.1 | Weak |
| sdc-2 | Dosage compensation | No Hits | – | – | – | NP_509924.1 | Missing |
| sdc-3 | Dosage compensation | No Hits | – | – | – | NP_506703.1 | Missing |
| **Somatic Sex Determination** | | | | | | | |
| her-1 | Secreted masculinizer | Scaffold 6 | 1.3 | 26.9 | – | NP_001024310.1 | Missing |
| tra-2 | Membrane receptor | Scaffold 2 | 0.001 | 44.6 | – | NP_495426.1 | Missing |
| fem-3 | Intracellular signaling | Scaffold 4 | 2.3 | 31.4 | – | NP_001255365.1 | Missing |
| **Signaling & Adapters** | | | | | | | |
| fem-1 | Intracellular signaling | Scaffold 2 | 2.30e-27 | 70.0 | OG0002917 | NP_500824.1 | Present |
| fem-2 | Intracellular signaling | Scaffold 1 | 7.35e-21 | 94.7 | OG0005340 | NP_497224.1 | Present |
| cul-2 | Ligase Scaffold | Scaffold 1 | 7.70e-24 | 108 | OG0001067 | NP_001023008.1 | Present |
| elc-1 | Ligase Adaptor | Scaffold 1 | 5.98e-34 | 113 | OG0001432 | NP_001380023.1 | Present |
| zyg-11 | Cell Cycle / Meiosis | Scaffold 4 | 4.03e-12 | 70.8 | – | NP_495677.1 | Weak |
| **Somatic Sex Determination Regulator** | | | | | | | |
| tra-1 | Regulation of Sex Determination | Scaffold 1 | 4.78e-48 | 99.5 | OG0001538 | NP_001022880.2 | Present |
| **Germline Effectors** | | | | | | | |
| fog-2 | Germline switch | Scaffold 3 | 3.1 | 30.3 | – | NP_001041187.1 | Missing |
| gld-1 | Germline repressor | Scaffold 2 | 7.87e-28 | 115 | – | NP_492143.1 | Weak |
| fbf-1/2 | Germline fate | Scaffold 4 | 1.30e-46 | 141 | – | NP_495216.1 | Weak |
| fog-1 | Germline switch | Scaffold 4 | 5.24e-07 | 53.6 | – | NP_001367957.1 | Weak |
| fog-3 | Germline switch | Scaffold 5 | 2.28e-05 | 45.2 | OG0007206 | NP_492646.1 | Weak |
| **Somatic Male Targets** | | | | | | | |
| mab-3 | Somatic Male Switch | Scaffold 2 | 1.84e-13 | 69.7 | OG0003005 | NP_001022464.1 | Present |
| ceh-30 | Male Neuron Survival | Scaffold 1 | 4.73e-27 | 107 | OG0002721 | NP_508524.2 | Present |
| egl-1 | Apoptosis Activator | No Hits | – | – | – | NP_506575.2 | Missing |
| mab-23 | Male Ray Development | Scaffold 2 | 0.51 | 27.8 | – | NP_001041090.1 | Missing |
| plx-2 | Axon Guidance | Scaffold 4 | 5.73e-22 | 70.5 | – | NP_001364609.1 | Weak |
| **Regulatory Partners** | | | | | | | |
| daf-16 | Aging / Diapause | Scaffold 5 | 7.53e-32 | 129 | – | NP_001367893.1 | Weak |
| hen-1 | Nerve ring signaling | Scaffold 5 | 1.54e-05 | 42.1 | – | NP_509200.1 | Missing |
